# Generation of a *Plasmodium falciparum* reporter line for studies of parasite biology throughout the life cycle

**DOI:** 10.1101/2023.05.23.542002

**Authors:** Pablo Suárez-Cortés, Giulia Costa, Manuela Andres, Daniel Eyermann, Cornelia Kreschel, Liane Spohr, Christian Goosmann, Volker Brinkmann, Elena A. Levashina

## Abstract

Fluorescence reporter strains of human malaria parasites are powerful tools to study the interaction of the parasites with both human and mosquito hosts. However, low fluorescence intensity in transmission-relevant parasite stages and the choice of insertion loci that cause parasite developmental defects in the mosquito largely limits usefulness of currently available lines. To overcome these limitations, we used a CRISPR-Cas9-mediated approach to generate *PfOBC13^GFP^*, a novel selection marker-free reporter parasite in the background of the African NF54 *Plasmodium falciparum* line. As docking site, we selected the *OBC13* locus that is dispensable for asexual and sexual development *in vitro*. *PfOBC13^GFP^* parasites encode GFP flanked by *hsp70* UTRs that drive strong fluorescence reporter expression throughout blood and mosquito stages, enabling parasite detection by such high throughput methods as flow cytometry. When compared to the parental line, *PfOBC13^GFP^* parasites showed normal development during blood and mosquito stages, and they efficiently infected the main African vector *Anopheles coluzzii,* overcoming one of the limitations of the previously developed fluorescent reporter lines based on the *Pfs47* locus. *PfOBC13^GFP^* constitutes a potent tool enabling host-pathogen studies throughout *Plasmodium* life cycle.

**Importance:** Fluorescence reporter strains have been very useful in malaria research, however, up to date they had limitations in mosquito infectivity and fluorescence intensity. Here we report the generation of *PfOBC13^GFP^*, a new fluorescent parasite strain of the human malaria parasite *P. falciparum*. *PfOBC13^GFP^* parasites are highly fluorescent throughout the life cycle, making them an ideal tool for the study the parasite progression through blood and mosquito stages. They efficiently infect the African mosquito *vector A. coluzzii*, allowing the study of this African parasite in its biological background. Moreover, strong parasite fluorescence enables flow cytometry and live microscopy characterization of all parasite stages, especially those involved in transmission.

## Introduction

*Plasmodium falciparum* is the deadliest human malaria parasite, with a current estimated annual death toll above 600,000 (1). To develop new strategies for disease control and prevention, it is essential to get a better understanding of the parasite biology throughout its complex life cycle that alternates between mammalian host and mosquito vector.

Transgenic malaria parasites expressing reporter proteins have proven to be powerful tools to better understand parasite-host interaction (2) or establish *in vitro* high throughput growth inhibition screens targeting asexual stages (3) and gametocytes (4). The first fluorescence malaria parasites reporter lines generated for rodent malaria parasites greatly improved our knowledge of the parasités life cycle and biology (5). However, some aspects of the biology of these model parasites markedly diverge from the human malaria parasites (6), highlighting the need for *P. falciparum* fluorescence reporter lines to study the biology of human malaria. Advances in transgenesis benefited establishment of *P. falciparum* fluorescence reporter lines, first by episomal material (3) and later by single homologous recombination techniques, both of which, however, have very low efficiency in *P. falciparum* and result in non-stable transgenic lines that must be kept under drug selection to ensure retention of the reporter cassette (7, 8). Recent introduction of CRISPR/Cas9-based techniques (9) greatly improved the efficiency of parasite transgenesis, leading to the faster generation of stable, selection marker-free fluorescence reporter lines (10-12). However, selection of stage-specific promoters and lack of information about insertion loci resulted in variable levels of expression of the reporter gene during parasite life cycle and low mosquito infectivity of the reporter lines.

Because of the scarcity of well characterized loci, previously generated *P. falciparum* fluorescence reporter lines were based on stable integration of the expression cassette into *Pfp230p* or *Pf47* loci (7, 10-14). As both genes were essential for *P. falciparum* reproduction, their disruption prevented studies that targeted mosquito stages of parasite development and transmission (15, 16). As a result, the reporter lines with insertions in these genes have been mostly used to study sexual blood stages and *P. falciparum* development in the Asian malaria vector *A. stephensi* (7, 13, 14) but not in the main African vector *A. gambiae* s.l. Very recently, a new fluorescence reporter line was generated by insertion into the *pfpare* locus, however so far this line was only used for *in vitro* characterization and the information about its mosquito-infecting capacity is still missing (17).

Another issue of the currently used fluorescence reporter lines is low fluorescence levels in some parasite stages, including the infective to humans sporozoites. Sporozoites are the target of the only licenced malaria vaccine. A *P. falciparum* line with strong fluorescence reporter expression in sporozoites will benefit flow cytometry and *in vivo* imaging analyses that are instrumental for immunological and functional assays in the context of vaccine development. So far, the promoters designed to drive strong expression of fluorescence reporters either target specific stages (3, 11, 14, 18, 19), or multiple stages by the use of promoters of such housekeeping genes as *sui1* (12), *40s* (12), *cam* (10), *gadph* (10), *ef1α* (7, 8, 11, 12, 14) or *hsp70* (10, 13, 17). Although *ef1α* has been frequently used in *Plasmodium* reporter strains, expression of fluorescence markers under this promoter is weak in sporozoites, thereby limiting the use of this promoter in transmission studies (14). Currently, *hsp70* stands out as a potent constitutive promoter for reporter expression throughout the whole life cycle, first in the in *P. berghei* (20) and more recently in strong multistage reporter expression in *P. falciparum* (10, 13, 17).

Here we report the generation of a selection marker-free stable transgenic *P. falciparum* parasite line *PfOBC13^GF^* that shows strong *hsp70*-driven GFP expression throughout asexual, sexual and transmission stages. The reporter cassette was inserted into a new *obc13* locus that was reported to be expressed in female gametocytes but whose function was not required for gamete egress and round up *in vitro* (21). We demonstrate that *PfOBC13^GFP^* parasites efficiently infect both African and Asian mosquito vectors *Anopheles coluzzii* and *A. stephensi*, respectively. Systematic comparison of *PfOBC13^GFP^* development and functions with the parental line, identified an unexpected role of OBC13 *in vivo* as disruption of this *locus* resulted in a two-fold decrease in the number of developed ookinetes in the mosquito midgut. However, this defect in ookinete development did not significantly affect further parasite development in the vector as the number of generated sporozoites and their infectivity to hepatocytes *in vitro* were similar between *PfOBC13^GFP^*and the parental line. Our results suggest that the *PfOBC13^GFP^*line constitutes valuable tool to study the biology of mosquito-parasite interactions and parasite transmission blocking applications.

## Materials and Methods

### P. falciparum culture

*P. falciparum* (NF54 line (22)) was cultured in O^+^ human erythrocytes (Haema, Berlin), in a 4% CO_2_, 3% O_2_ atmosphere, at 37°C. NF54 line originated at Prof. Sauerweins’s laboratory at the Radboud University Medical Centre, Nijmegen, The Netherlands, and was monthly tested for mycoplasma contamination. Asexual parasites were kept at 3-4 % haematocrit in RPMI 1640 medium with L-glutamine, 25 mM HEPES, 10 mM hypoxanthine, 20 mg/ml gentamicin, supplemented with 10% A^+^ human serum. Gametocytogenesis was induced by setting asynchronous asexual cultures at 4% haematocrit and 1% parasitaemia, with complete medium without gentamycin replaced every day until D14 post seeding. Stage V gametocytes were then quantified microscopically using a cell counting chamber (Neubauer) and the day after they were used for mosquito infections. Alternatively, for *PfOBC13KO*, *PfOBC13^GFP^* and NF54 comparisons (Fig. S4), gametocytogenesis was induced by setting asynchronous asexual cultures at variable parasitaemias to reach 4-5% on day 1 post seeding, and complete medium without gentamycin replaced every day except on day 2 post-seeding, when 50% of the spent medium was kept. On day 12 post-seeding, stage V gametocytes were quantified microscopically using a cell counting chamber (Neubauer) and the day after they were used for mosquito infections.

### Generation of transgenic parasites

Plasmids *pOBC13^GFP^* and *pCBS-OBC13* (see Supplementary methods) were transfected as described elsewhere (23). Briefly, 200 μl of packed RBCs containing young ring stage parasites at 3-4% parasitaemia were electroporated in the presence of a 1:1 mixture of plasmids (200 μg of DNA in total diluted in 400 μl of Cytomix). Transfected parasites were cultured for 24 h in 5 ml of fresh media with 5% haematocrit without drug selection. From day 2 to day 7 post-transfection, medium was supplemented with 5 μg/ml blasticidine S (Cayman) for parasite selection. Medium was changed every day for the first 7 days and every 3 days after that. Fresh 50 μl of packed red blood cells were added every 7 days until parasites were detected in thin blood smears after 25 days. Parasites were cloned by serial dilution to obtain clonal strains and tested for transgene integration by PCR. The obtained line was named *PfOBC13^GFP^*.

### Mosquito rearing and parasite infections

*Anopheles coluzzii* mosquitoes (Ngousso strain) were maintained at 28°C and 70-80% humidity with a 12/12 day/night cycle. For parasite infections, stage V gametocytes were mixed with fresh red blood cells and human serum (50% haematocrit) to a concentration of approximately 3.7x10^6^ gametocytes/ml. Mosquitoes were fed for 15 min on a membrane feeder, and unfed mosquitoes were removed shortly after feeding. For sporozoite generation, mosquitoes were given an additional blood meal 7 dpi (days post infection). Infected mosquitoes were kept in a BSL3 laboratory at 26°C for up to 15 days according to national regulations (Landeamt für Gesundheit und Soziales, project number 297/13).

### Characterization of asexual and sexual stages *in vitro*

Replication rate of asexual cultures was calculated as the increase in parasitaemia over 48 h taking into account the dilution factor at the first time point.

Sexual commitment was calculated by counting gametocytaemia on day 7 after start of gametocytogenesis. Cultures were smeared, stained with Giemsa and gametocytes (stage II to V) and RBCs were counted under a microscope.

Exflagellation rates of gametocyte cultures were quantified after induction of 200 μl of culture with 20 μM of xanthurenic acid (XA, Sigma,) for 15 min at 26°C. Exflagellation centres were enumerated microscopically in a counting chamber (Neubauer). Exflagellation rate was calculated as the number of exflagellation centres per ml divided by the number of stage V gametocytes per ml.

Gametocyte activation (round-up and egress) was measured as previously described (24). Briefly, at day 14 post-induction gametocyte cultures were Percoll-purified, resuspended in RPMI 1640 medium and stained for 5 min with 1 μg/ml of WGA-CF594 (Biotium). Gametogenesis was induced by XA at the concentration indicated for each experiment, ranging from 0.1 to 100 μM. Gametogenesis was stopped 15 min after initiation by addition of 1% paraformaldehyde (PFA) at RT. After parasite centrifugation, cells were resuspended in 10 μl PBS and mounted on a microscope slide. 200 sexual parasites were identified using bright field microscopy and classified by morphology as gametocytes (falciform) or gametes (round). Egress was evaluated by the presence or absence of WGA signal fluorescence surrounding the parasites for 200 gametes.

### Mosquito dissections

Mosquitoes were decapitated and their midguts dissected at the indicated time points (20 midguts per sample). For oocyst count and observation, mosquitoes were killed in ethanol 70% at 8 to 11 dpi and then rinsed in PBS. For experiments involving sporozoites, mosquitoes were killed in 70% ethanol at 13 to 15 dpi and the salivary glands from 40 to 100 mosquitoes were dissected in complete HC-04 medium and homogenized using a glass homogenizer. Sample was filtered through a 40 μm strainer and sporozoites were counted on a cell counting chamber (Malassez, Marienfelde). Sample was kept on ice until further use.

### Ookinete staging and counting

Midguts of 20 mosquitoes were pooled and homogenized in 200 μl PBS containing Hoechst (1:500, Molecular Probes) and anti-Pfs25 AlexaFluor568-conjugated antibody (7.5 and 1.5 μg/ml at 2 and 24 h post infection (hpi), respectively) (25). After incubation at 4°C for 30 min, samples were washed with 1.5 ml PBS and pelleted for 4 min at 300 g on a table top centrifuge. Cells were resuspended in 100 μl PBS and Pfs25-positive cells were counted and classified as described previously (25).

### Oocyst staining and counting

Mosquito midguts were dissected 8-11 days post infection in PBS and stained for 10 min with 1% mercurochrome in water in a humidified chamber. Oocysts were counted for each midgut under a light microscope (30 midguts per condition).

### Sporozoite gliding assay

Hybridoma cells expressing anti-PfCSP IgG (clone 2A10) were obtained from BEI Resources. IgG fraction was purified using Protein G Sepharose columns (GE Healthcare) and part of it was labelled with AlexaFluor568 (Life Technologies) in the Central Laboratory Facility (Deutsches Rheuma-Forschungszentrum, Berlin). Sporozoite gliding was performed using previously described methods (26). In brief, glass Lab-Tek 8-wells (Nalgene) were coated with 200 μl of 5 μg/ml anti-PfCSP mAb (2A10 clone) in PBS overnight at RT and washed with PBS. Sporozoites were diluted in HC-04 medium supplemented with 10% FCS to a concentration of 250 sporozoites/μl and 200 μl of this suspension was added to a well. After 1 h incubation at 37°C in 5% CO_2_, the medium was removed and the cells were fixed with 4% PFA in PBS, washed and blocked with 1% BSA in PBS. To visualize CSP trails, sporozoites were incubated for 2 h at RT with anti-PfCSP AlexaFluor568-conjugated mAb (clone 2A10, 0.836 μg/ml in PBS), washed and mounted in Aqua/Polymount medium (Polysciences). Sporozoites (n=200) and trails were quantified by an Axio Observer Z1 fluorescence microscope (ZEISS) and the proportion of gliding sporozoites was determined as the number of trails divided by the number of sporozoites.

### Sporozoite traversal assay

HC-04 cells (MRA-975, deposited by Jetsumon Sattabongkot (Sattabongkot et al., 2006) were cultured at 37°C at 5% CO_2_ using HC-04 complete culture medium (1:1 MEM (-L-glutamine)/F-12 Nutrient Mix (+ L-glutamine) media, 15 mM HEPES, 1.5 g/l NaHCO3, 2.5 mM L-glutamine and 10% FCS). Hepatocyte traversal assay was performed as previously described (27). In brief, salivary gland sporozoites (50,000 per well) were incubated with HC-04 cells (60,000 per well) for 2 h at 37°C at 5% CO_2_ in the presence of 0.5 mg/ml dextran-rhodamine (MW=10,000, Molecular Probes). To test sporozoite hepatocyte traversal in the absence or presence of anti-CSP mAbs, sporozoites were pre-incubated with 100 µg/ml of the non-PfCSP-reactive monoclonal antibody mGO53 (28) or a chimeric humanized version of the PfCSP-reactive monoclonal antibody 2A10 (27) for 30 min on ice. After sporozoite incubation, HC-04 cells were washed and fixed with 1% PFA in PBS before quantifying dextran-positive cells by flow cytometry (LSR II instrument, BD Biosciences). For data analysis using FlowJo V.10.0.8 (Tree Star), the background signal (dextran-positive cells treated with uninfected mosquito salivary gland material) was subtracted and the results were normalized to the maximum sporozoite traversal capacity (dextran-positive cells treated with salivary gland sporozoites without mAb addition).

### Live imaging of *P. falciparum*

*PfOBC13^GFP^* parasites were stained for 10-15 min with Hoechst (1:500, Molecular Probes) without washes or fixation. Z-stack projections (3 to 5 per parasite) were captured by an Axio Observer Z1 fluorescence microscope (ZEISS) using 40x and 63x objectives and Max Intensity Projections were generated and analysed by FIJI (ImageJ2 version 2.3.0/1.53q).

### Statistical Analysis

No samples were excluded from the analyses. Mosquitoes from the same batches were randomly allocated to the experimental groups (age range: 1–2 days). The experimenters were not blinded for the group allocation during the experiment and/or when assessing the outcome. Sample sizes were chosen according to best practices in the field and previous data (25). Statistical analyses were performed with GraphPad Prism 9 using tests indicated in the Figure legends. *P*-values below 0.05 were considered significant.

## Results

### Generation of the *P. falciparum* GFP omni-stage reporter line

To generate a parasite reporter line expressing fluorescence at all stages throughout the *P. falciparum* life cycle, we designed an expression cassette containing a codon-optimized version of Green Fluorescence Protein (GFP) adapted to the highly AT-skewed parasite genome. The reporter was flanked by the 5’- and 3’-UTRs of the constitutively expressed gene *Pfhsp70* (PF3D7_0818900). For the transgene insertion, we selected the *OBC13* (PF3D7_1214800) locus which was previously reported to be dispensable for asexual parasite growth, gametogenesis and gamete egress *in vitro* (21). To confirm that deletion of this gene did not impact overall parasite development in *A. coluzzii*, we first generated a new *PfOBC13KO* line in the *P. falciparum* NF54 background (Fig. S1 A and B). Evaluation of the *PfOBC13KO* infectivity to *A. coluzzii* mosquitoes did not reveal any differences in oocyst prevalence and loads between wild type (WT) and *PfOBC13KO* parasites (Fig. S1C). These results validated the use of the *OBC13* locus as a neutral insertion site for *P. falciparum* transgenesis.

*P. falciparum* parasites of the NF54 strain were transfected with the plasmids *pOBC13^GFP^* and *pCBS-OBC13*, that enabled the CRISPR/Cas9-mediated insertion of the GFP reporter cassette into the *OBC13* locus (Fig. S2 A). GFP-positive parasites were cloned by limiting dilution, and the correct transgene insertion into the *OBC13* locus was confirmed by diagnostic PCR (Fig S2 B). The resulting parasite line was named *PfOBC13^GFP^*.

### Characterization of *PfOBC13^GFP^* reporter line in asexual and sexual blood stages

We then examined GFP expression in live *PfOBC13^GFP^* parasites at all stages throughout the complex life cycle of *P. falciparum*. Asexual parasite stages of *PfOBC13^GFP^*, including rings, trophozoites, schizonts and merozoites showed a strong GFP signal (Fig. 1, upper-left panel) that was also detected in gametocytes and gametes after activation by xanthurenic acid *in vitro* (Fig. 1, lower-left panel).

**Figure 1.**
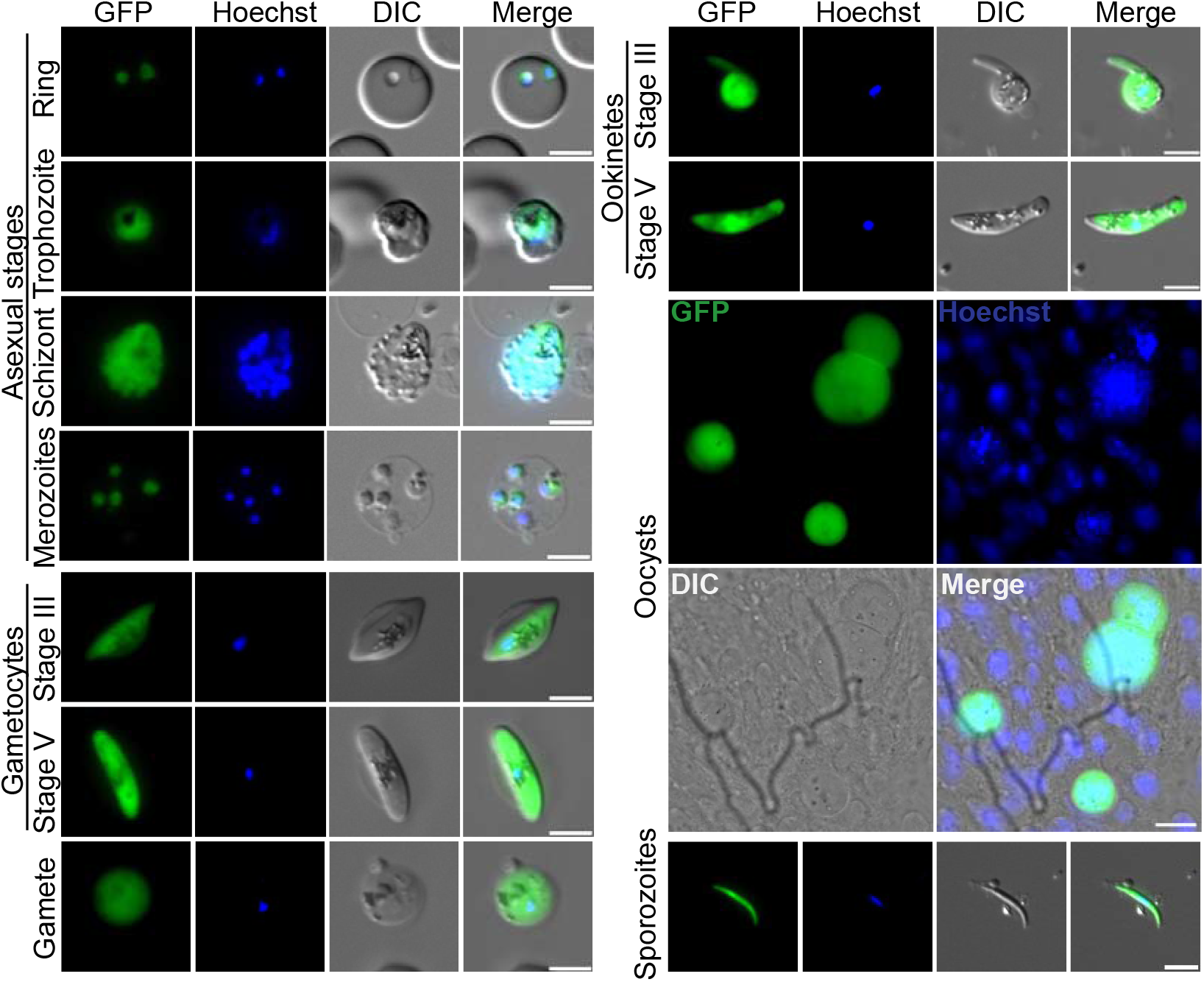
Expression of the fluorescence reporter in asexual, sexual and mosquito stages of *PfOBC13^GFP^*parasites. Fluorescence microscopy images showing live GFP signal of different blood and mosquito stages of *PfGFP^OBC13^* parasites. Asexual parasite samples from asynchronous cultures, sexual stages from gametocyte cultures at day 7 (stage III) and day 14 post induction (stage V and gamete). Ookinetes were obtained from blood bolus of infected mosquitoes 24 hpi. Oocysts observed from a dissected mosquito midgut 11 dpi. Sporozoites were obtained from salivary glands of mosquitoes 15 dpi. Scale bars 20 μM (oocysts panel) and 5 μM (all other panels).

To evaluate reporter expression in the mosquito stages, *A. coluzzii* mosquitoes were fed with *PfOBC13^GFP^* gametocyte cultures and their midguts were dissected at multiple time points after infection. GFP signal was observed in ookinetes, oocysts and sporozoites (Fig. 1, right panel). These results demonstrated that the transgenic *P. falciparum* parasite expressed the fluorescence reporter cassette at all examined stages.

In order to evaluate the fitness of the newly generated *PfOBC13^GFP^* line, we next compared its asexual growth rates and gametocyte production capacity with the parental NF54 (WT) line. Asexual growth rate was evaluated as fold-change in parasitaemia of asynchronous cultures in 48 h, and gametocyte production as the number of maturing gametocytes at day 7 post induction of gametocytogenesis. No differences were observed for these parameters between WT and *PfOBC13^GFP^*transgenic parasites (Fig. 2 A, B).

**Figure 2.**
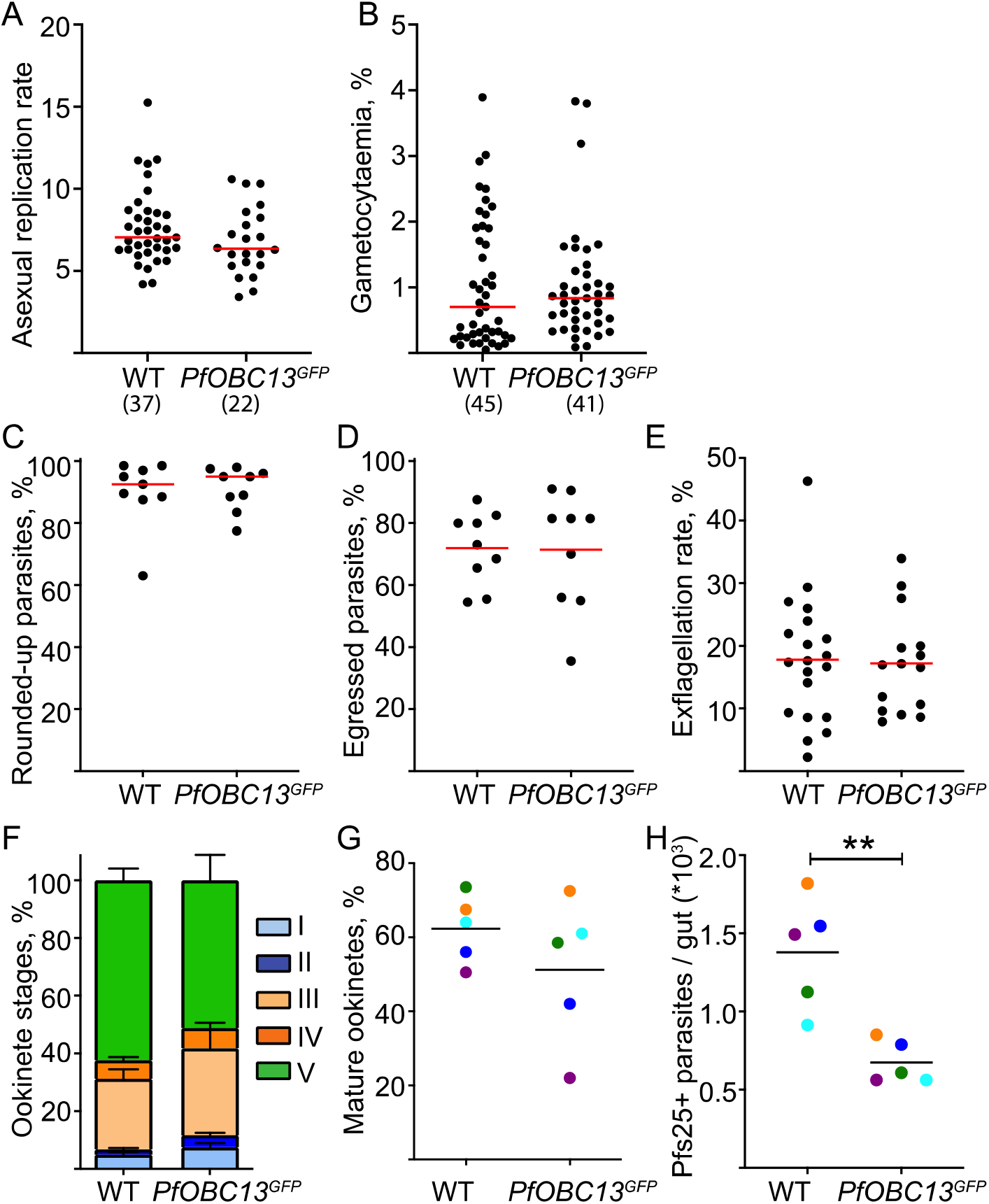
Characterization of *PfOBC13^GFP^* blood and early transmission stages *in vitro* and *in vivo*. (**A**) Asexual replication rate was calculated for WT and *PfOBC13^GFP^* parasites as the fold change in parasitaemia in 48h in asynchronous asexual cultures. Unpaired *t*-tests were performed to compare samples (N = 37 (WT), N = 22 (*PfOBC13^GFP^*), mean shown). (**B**) Gametocytaemia in WT and *PfOBC13^GFP^* cultures was measured at day 7 post-induction, counting gametocytes (stages II to IV) on Giemsa-stained thin smears. Unpaired *t*-tests were performed to compare samples (N = 45 (WT), N = 41 (*PfOBC13^GFP^*), mean shown). Gametogenesis of WT and *PfOBC13^GFP^* parasites were examined in parallel after activation with xanthurenic acid (XA) *in vitro*. (**C**) Gametocyte round-up rate (calculated as percentage to total number of sexual parasites) and (**D**) Egress rate (calculated as percentage of red blood cell-free round gametes to the total number of rounded-up gametocytes) in culture 15 min after activation with XA (100 μM). Each dot indicates mean of one gametocyte culture. Statistically significant differences were evaluated by Unpaired *t*-test (N = 9, n = 200, mean shown). (**E**) Exflagellation rate (proportion of exflagellation clusters of total number of stage V gametocytes) 15 min post activation with XA (20 μM). Each dot indicates mean of one gametocyte culture. Statistically significant differences were evaluated by Unpaired *t*-test (N = 20 (WT), N = 15 (*PfOBC13^GFP^*), mean shown). Ookinete development in the blood bolus of infected mosquitoes at 24 hpi *in vivo*: ookinete stages I to V (**F**) or stage V alone (**G**) are indicated from the same experiments. Statistically significant differences were evaluated by Wilcoxon test (N = 5, n = 200, error bars indicate SEM, mean shown). (**H**) Number of Pfs25-positive (Pfs25^+^) parasites per midgut at 24 hpi. Statistically significant differences were evaluated by Paired *t*-test (N = 5, n = 200, mean shown). Color code shows paired experiments. **\*\***: p < 0.01, nonsignificant differences are not indicated.

Next, we addressed the efficiency of gamete formation in the reporter parasites. Gametogenesis is triggered by changes by the environmental cues in the mosquito midgut upon ingestion of gametocytes leading to the parasite rounding up and egress from the red blood cells. In addition, male gametes undergo triple nuclear division to generate flagellated gametes that rapidly propel themselves in search of a female gamete in a process called exflagellation. To evaluate round-up, egress and exflagellation rates of *PfOBC13^GFP^* gametocytes *in vitro*, we first determined optimal activating XA concentrations in WT parasites (Fig. S3). The optimal concentration of XA for gametocyte round-up and egress was observed at 100 μM, whereas the highest exflagellation rates were detected at 20 μM. Comparison of WT and *PfOBC13^GFP^* showed no differences (Fig. 2 C-E), supporting our earlier observations that disruption of the *OBC13* locus did not impact gametocyte activation *in vitro* (21).

### Characterization of *PfOBC13^GFP^* development in *A. coluzzii*

To evaluate development of the reporter line *in vivo*, *A. coluzzii* mosquitoes were infected with WT and *PfOBC13^GFP^*gametocyte cultures. Mosquito midguts were dissected 24 h post infection (hpi), and the early infection stages of parasites were microscopically examined in the blood bolus. Extracellular parasites at this stage include unfertilized round egressed female gametes zygotes and developing ookinetes, ranging from round-shaped (stage I) to mature banana-shaped (stage V) (25). Because all these stages express the surface protein Pfs25 (25), we used it as a marker to visualize parasite development in WT and *PfOBC13^GFP^* lines by immunofluorescence analyses. No differences were observed in the relative proportions of ookinete stages at 24 hpi, suggesting normal ookinete maturation in the *PfOBC13^GFP^*line (Fig. 2 F, G). However, a two-fold decrease in overall parasite numbers was detected in *PfOBC13^GFP^* parasites compared to WT (Fig. 2 H). To explore this unexpected phenotype, we infected *A. coluzzii* mosquitoes with WT, *PfOBC13^GFP^*and *PfOBC13KO* parasites in parallel. Quantification of Pfs25-positive parasites confirmed lower parasite numbers in Pf*OBC13^GFP^*and *PfOBC13KO* lines compared to WT, both at early (2 hpi) and late (24 hpi) time points of ookinete development, while ookinete maturation rates remained similar (Fig. S4 A, B). These results suggested that disruption of the *PfOBC13 locus* had a mild negative effect on gametogenesis *in vivo*. *OBC13* is expressed in osmiophilic bodies, an organelle of female gametocytes with a demonstrated role in gametogenesis (21). As some proteins of osmiophilic bodies have been shown to be essential for the formation of these organelles (29), we examined the morphology of osmiophilic bodies in *PfOBC13^GFP^* and *PfOBC13KO* gametocytes by transmission electron microscopy. We found that *OBC13* disruption did not impact osmiophilic body formation (Fig S5), suggesting that the observed defect in gametogenesis *in vivo* was not linked to biogenesis of osmiophilic bodies.

We next examined oocyst formation in infected mosquitoes by dissecting mosquito midguts at 11 days post infection (dpi). Despite the high variability in oocyst yields between experiments, we found that both the proportion of infected midguts (prevalence of infection) and the oocyst numbers per midgut (intensity of infection) were similar between WT and *PfOBC13^GFP^* parasites (Fig. 3 A, B). This data confirmed our previous results with the *PfOBC13KO* line (Fig. S1 C).

**Figure 3.**
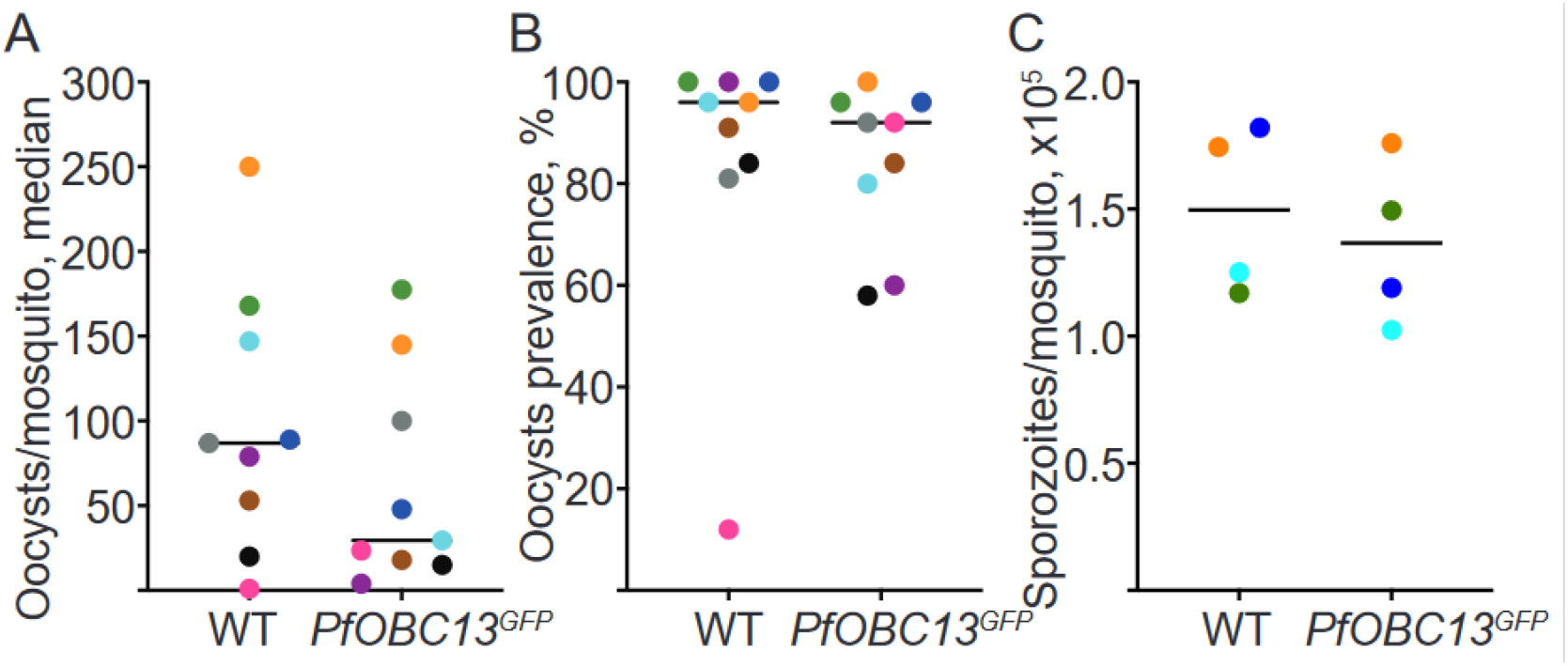
Characterization of oocyst and sporozoite stages of the *PfOBC13^GFP^* line. WT and *PfOBC13^GFP^* oocyst infection (**A**) intensity (median) and (**B**) prevalence (%) in *A. coluzzii* mosquitoes. Statistically significant differences were evaluated by Wilcoxon test (N = 9, n = 13-30, median shown). (**C**) Mean sporozoites per mosquito, counted after pooling of salivary glands isolated from 40-100 mosquitoes per condition 13-15 dpi. Statistically significant differences were evaluated by Paired *t*-test (N = 4, mean shown). Color code indicates paired experiments. Nonsignificant differences are not indicated.

We then quantified sporozoite production. Similar sporozoite numbers per mosquito were observed for WT and *PfOBC13^GFP^* parasites at 13-15 dpi (Fig. 3 C). Similar results were observed in infections of the Asian mosquito vector *A. stephensi* (Fig. S6). In conclusion, despite mild phenotype in gametogenesis, the *PfOBC13^GFP^* line was as efficient in infection of *Anopheles* mosquitoes as the parental line.

### Motility and infectivity of *PfOBC13^GFP^* sporozoites

We next examined sporozoite motility and infectivity of the fluorescent reporter line. Both gliding motility on a glass slide and the capacity to traverse hepatocytes *in vitro* were similar between WT and *PfOBC13^GFP^* parasites (Fig. 4 A, B). The circumsporozoite protein (CSP) is the main surface antigen of sporozoites and the target of the only licenced malaria vaccine. In order to validate *PfOBC13^GFP^* sporozoites as a tool for functional studies of anti-CSP monoclonal antibodies (mAbs) and CSP-based immunogens, we examined CSP expression levels and function. Surface *CSP* expression levels in *PfOBC13^GFP^*sporozoites measured by immunofluorescence microscopy was similar between WT and *PfOBC13^GFP^* sporozoites (Fig. 4 C). Importantly, endogenous expression of GFP and CSP was also amenable for flow cytometry quantification (Fig. S7). Finally, we tested whether *PfOBC13^GFP^* sporozoites can be used as a tool to evaluate inhibitory capacity of anti-CSP mAbs (30). Incubation with the anti-repeat CSP antibody 2A10 (27) showed similar levels of traversal inhibition for WT and *PfOBC13^GFP^* sporozoites (Fig. 4 D). These results validated the potential of the new *P. falciparum* reporter line for CSP-based sporozoite functional studies by fluorescence-activated cell sorting (FACS).

**Figure 4.**
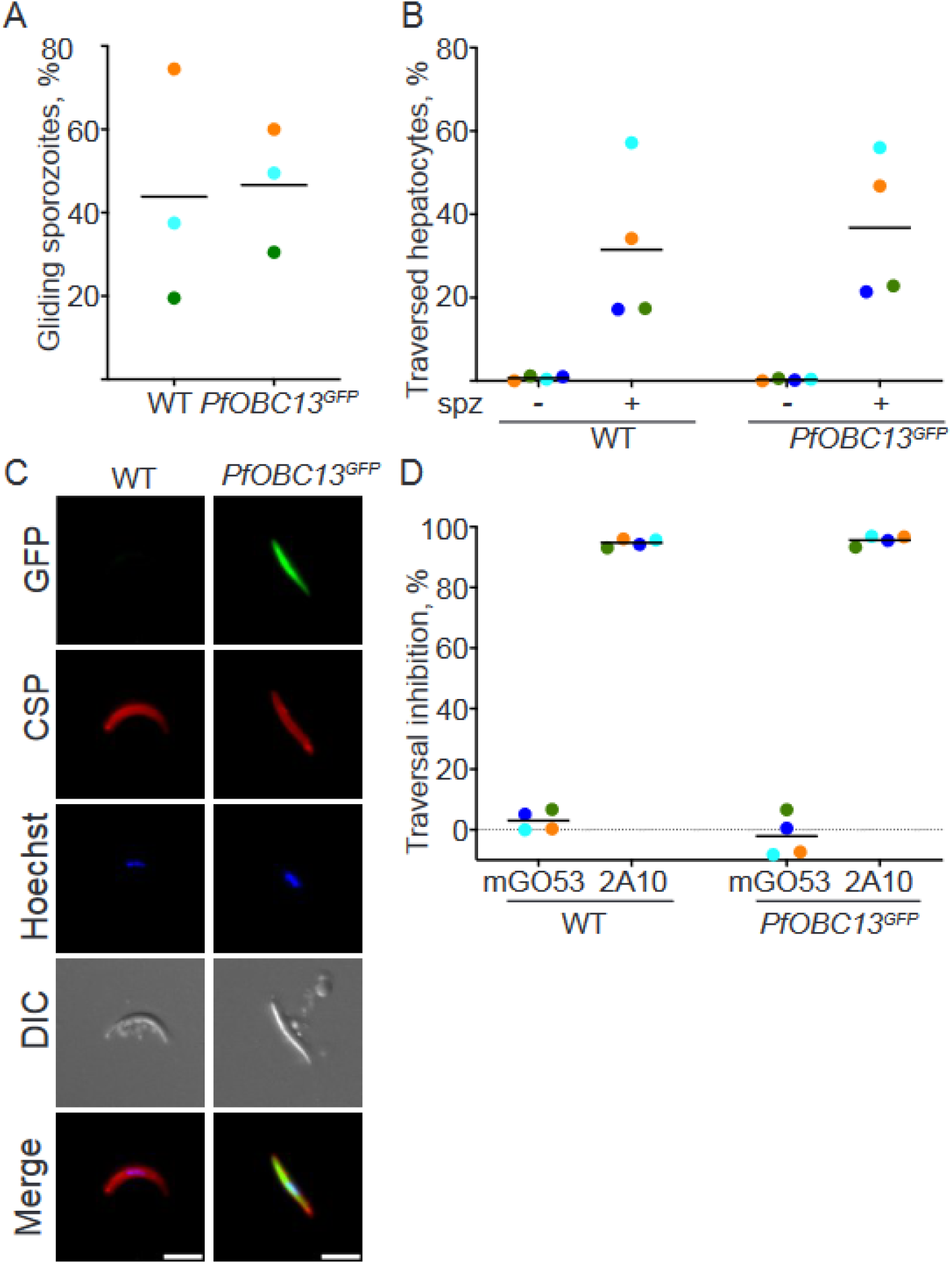
*PfOBC13^GFP^* sporozoites are functional and express normal levels of CSP. (**A**) Gliding capacity of isolated sporozoites on anti-CSP antibody coated slides. Statistically significant differences were evaluated by paired *t*-test (N = 3, n = 200, mean shown). (**B**) Hepatocyte traversal by the salivary gland sporozoites (spz). Material from uninfected salivary glands was used as control. Statistically-significant differences were evaluated by Friedman test (N = 4, mean shown). (**C**) Immunofluorescence analyses of the surfaceexpression of CSP in the wild type and *PfPOBC13^GFP^* transgenic sporozoites using α-CSP mAb (2A10 clone). Scale bar 5 μM. (**D**) Hepatocyte traversal inhibition of wild type and *PfOBC13^GFP^*sporozoites by anti-CSP mAb 2A10 and the isotype control mGO53 antibodies. Statistically significant differences were evaluated by Friedman test followed by Dunn’s multiple comparisons test (N = 4, mean shown). Color code indicates paired experiments. Nonsignificant differences are not indicated.

## Discussion

Fluorescence genome-encoded reporter lines of malaria parasites are useful tools to study parasite biology and host-parasite interactions. However, the majority of these reporter lines have been established for *Plasmodium* blood stages. Here, we developed a human malaria parasite line with strong constitutive GFP expression throughout *P. falciparum* blood and mosquito stages. Moreover, the absence of a drug selection marker in *PfOBC13^GFP^* should facilitate further genetic modification of the reporter line, for example knock-in or knockout modifications.

One of the shortcomings of the reported fluorescence lines is uneven fluorescence intensity in different parasite stages caused by selection of stage-specific regulatory sequences (7, 8, 11, 12, 14). To improve on that, we exploited 5’ and 3’ regulatory regions of *PfHsp70* gene, which has been reported to be strongly expressed throughout the entire *P. falciparum* life cycle (31, 32). Thus, an important advantage of *PfOBC13^GFP^* parasites is strong fluorescence that can be quantified by microscopy and flow cytometry.

Given the central role of CSP in malaria vaccine development, testing binding capacities and inhibitory activities of anti-CSP mAbs is key to improve current vaccine strategies (33). Previously reported *P. falciparum* reporter lines could in principle be useful for this purpose (11-14). However, these lines were generated by disrupting an important *Pfs47* locus rendering these lines unable to infect the African vector *A. coluzzii* (16). Therefore, reported here *PfOBC13^GFP^* reporter line can be used to study development of Africa-originated parasites in African mosquito vector.

In an attempt to find a neutral locus for transgene insertion into *P. falciparum* genome, we initially selected the *cg6 locus*, which has been shown to be dispensable for asexual and gametocyte parasite development *in vitro* and has been previously used as a docking site (34, 35). However, we found out that disruption of *cg6* caused a significant reduction in oocyst loads (data not shown).

The *obc13* locus has also been reported to be dispensable for asexual and sexual parasite development *in vitro* (21). However, infections in mosquitoes showed a two-fold decrease in the numbers of egressed parasites in *OBC13KO* and *PfOBC13^GFP^*lines compared to WT. Taken together, these results demonstrated that *in vivo* infections are more restrictive for mutant parasite lines and represent a more sensitive background for functional gene analyses compared to *in vitro* conditions.

Further characterization of *PfOBC13^GFP^* line uncovered that the observed decrease in the numbers of egressed parasites did not translate into lower levels of oocyst or sporozoite loads. As maximum 50 % of all ingested gametes form zygotes, it is possible that the impact of *OBC13* disruption on gamete numbers was minimized by fertilization efficiency. Alternatively, lower numbers of developed ookinetes at low parasite densities were more efficient in crossing the mosquito midgut epithelium and developing into productive oocysts. Further studies are needed to examine these hypotheses. Importantly for this study, our results showed that *obc1*3 plays a minor role at early mosquito stages and does not impact parasite infectivity to the vector. Although the identification of additional docking loci for *P. falciparum* would be beneficial, the *OBC13* constitutes a useful target locus for studies of interactions between *P. falciparum* and *A. coluzzii*.

In conclusion, *PfOBC13^GFP^*reporter line constitutes a potent tool to study the biology of human malaria parasites and their interactions with human and mosquito hosts.

## Acknowledgements

This project was supported by the 2018 GCAM grant OPP1210564 from the Bill and Melinda Gates Foundation.

## Abbreviations

ORF: Open Reading Frame
RBC: Red Blood Cell
XA: Xanthurenic Acid
UTR: UnTranslated Region
hpi: hours post infection
dpi: days post infection
FACS: Fluorescence-Activated Cell Sorting

## Supplementary figures

**Figure S1.**
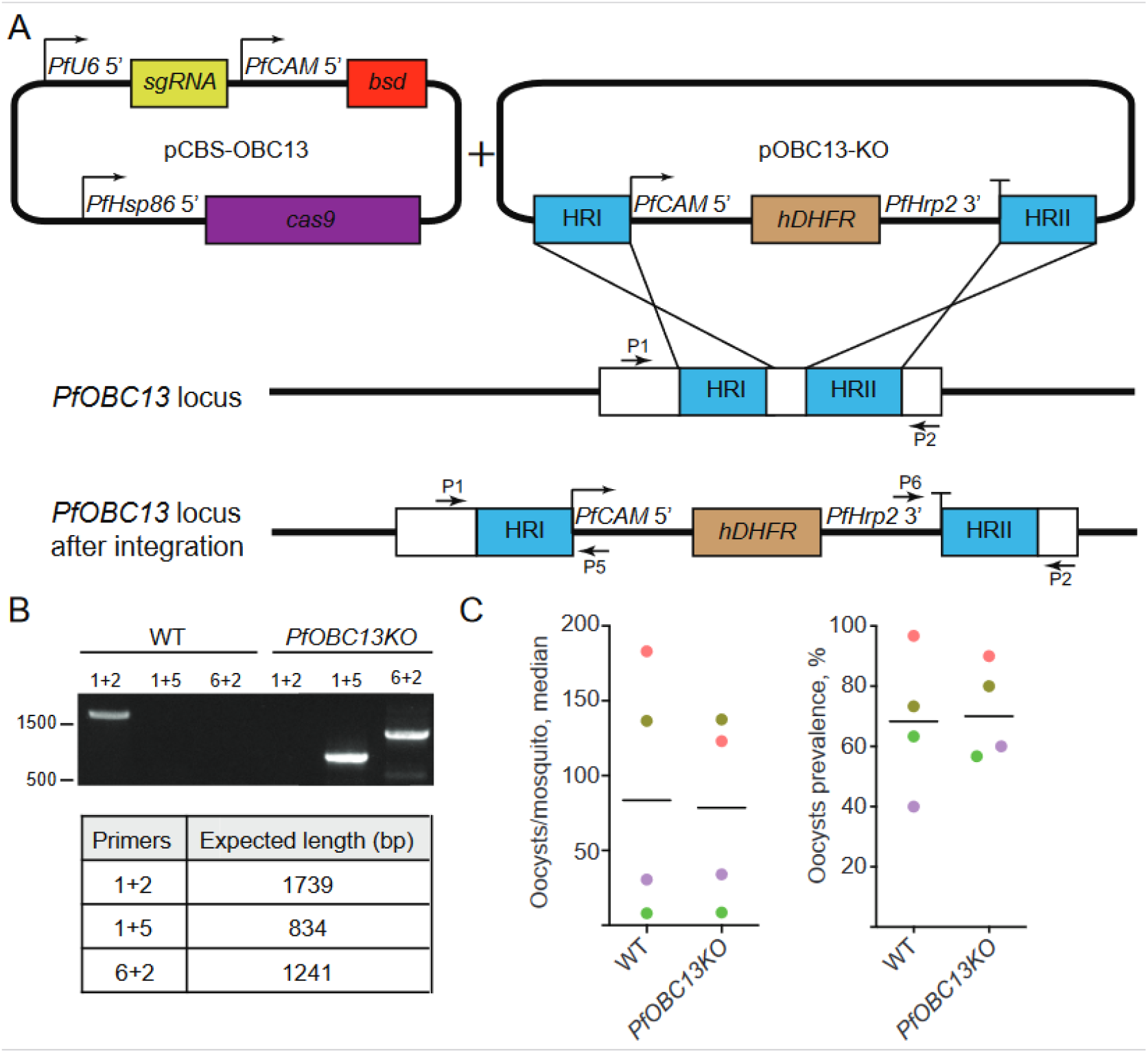
Generation and oocyst development of *PfOBC13KO* line. (**A**) Constructs used for the disruption of the *OBC13* locus in *P. falciparum* parasites. *pCBS-OBC13* was designed to target *OBC13* coding sequence, while *pOBC13-KO* provided a donor fragment which included the *hDHFR* expression cassette. Primers used in diagnostic PCRs are indicated by black arrows. PfU6: *Pf Ubiquitinase 6*, PfCAM: *Pf Calmodulin*, *PfHsp86*: *Pf Heat shock protein 86*, *PfHrp2*: *Pf Histidine rich protein 2*, *sgRNA*: single guide RNA, *bsd*: blasticidin resistance ORF, *cas9*: *cas9* ORF. *hDHFR*: human dihydrofolate reductase ORF. HR: Homology region. (**B**) Diagnostic PCRs showed complete substitution of the *OBC13^wt^* allele in transfected parasites. PCRs were performed on genomic DNA from WT and cloned *PfOBC13KO* parasites. (**C**) WT and *PfOBC13KO* oocyst infection intensity (mean oocysts per experiment) and prevalence (%) in *A. coluzzii* mosquitoes 9 dpi. Color code shows paired experiments. Paired Wilcoxon test was performed to compare samples (N = 4, n = 30, median shown). Nonsignificant differences are not indicated.

**Figure S2.**
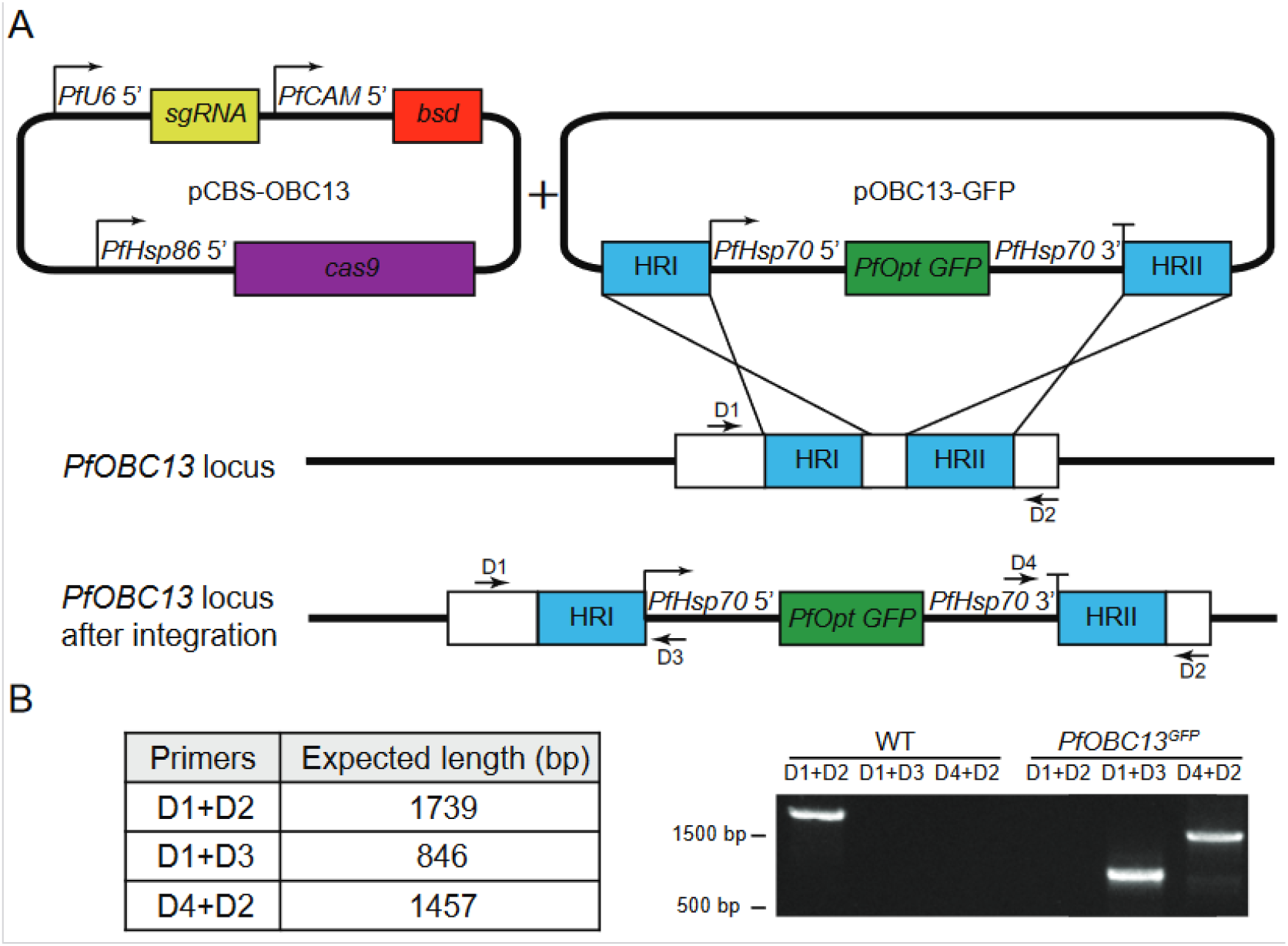
Generation of PfOBC13GFP line using CRISPR/Cas9 recombination. (**A**) Schematic representation of plasmids used for insertion of GFP expression cassette into the *OBC13* locus of *P. falciparum*. *pCBS-OBC13* targetsed the *OBC13* coding sequence, while *pOBC13-GFP* provided the donor GFP expression cassette. Primer positions (D1 to D4) used in diagnostic PCRs are indicated by black arrows. *PfU6*: *Pf Ubiquitinase 6*, *PfCAM*: *Pf Calmodulin*, *PfHsp86*: *Pf Heat shock protein* 86, *PfHsp70*: *Pf Heat shock protein 70*, *sgRNA*: single guide RNA, *bsd*: blasticidin resistance ORF, *cas9*: *cas9* ORF. *PfOpt GFP*: Codon-optimized GFP ORF. HR: Homology region. (**B**) Diagnostic PCRs confirmed the disruption of the *OBC13* locus and the presence of *PfHsp70* 5’ and 3’ UTRs in the transfected parasites. PCRs were performed on genomic DNA from WT and cloned *PfOBC13^GFP^* line. DNA and primer pair of each reaction are indicated.

**Figure S3.**
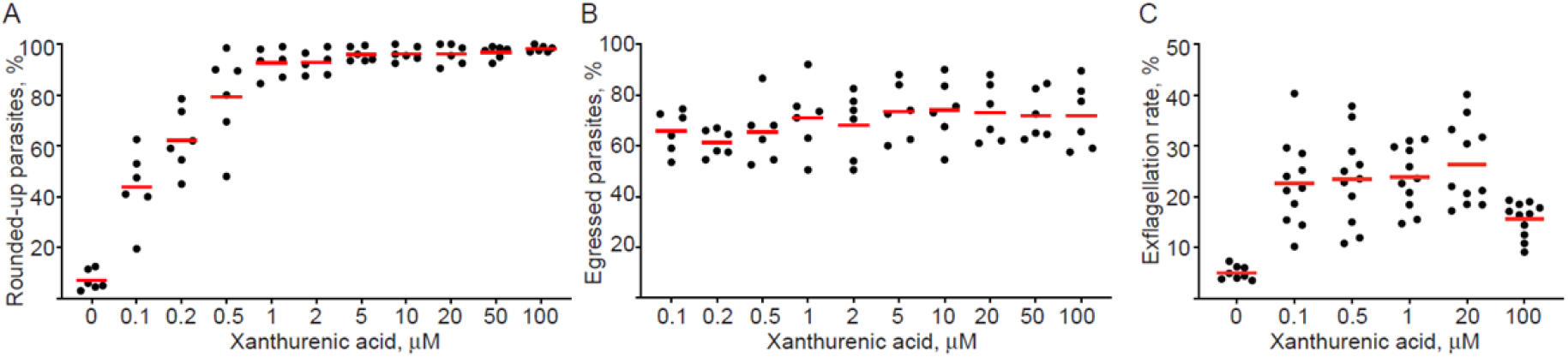
Dose-dependency of xanthurenic acid-induced gametocyte activation (rounding up, egress and exflagellation) in wild type parasites. WT NF54 mature gametocytes were activated with different concentrations of xanthurenic acid (XA). (**A**) Rounded-up parasites over total sexual parasites and (**B**) egressed gametes over total rounded-up parasites in culture 15 min post activation at the indicated XA concentrations are shown (N = 6, n = 200, mean shown). (**C**) Exflagellation rate (proportion of exflagellation clusters over total gametocytes stage V) 15 min post activation was determined at the indicated XA concentrations (N = 11, mean shown). Each dot indicates an independent gametocyte culture.

**Figure S4.**
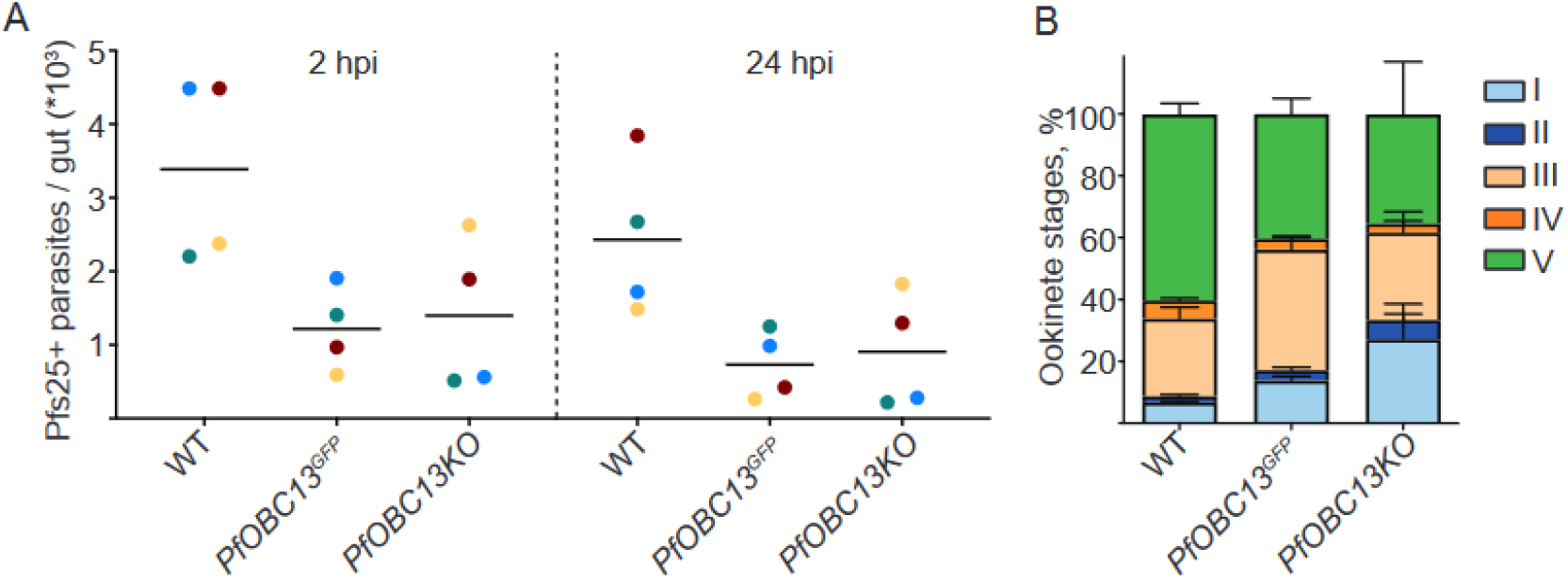
*C*haracterization of *PfOBC13^GFP^* and *PfOBC13KO* early transmission stages *in vivo*. *PfOBC13^GFP^*, *PfOBC13K*O and wild type (WT) parasites were examined in parallel to compare their phenotypes at the ookinete stage from blood bolus of infected mosquitoes. Number of Pfs25-positive (Pfs25*) parasites at 2 and 24 hpi (**A**) and proportions of ookinete stages I to V (**B**) are indicated from the same experiments (N = 4, n = 200, mean shown, error bars indicate SEM). Paired *Friedman* test corrected for multiple comparisons (Dunn’s test) was performed to compare samples in **A**. Color code shows paired experiments. Nonsignificant differences are not indicated.

**Figure S5.**
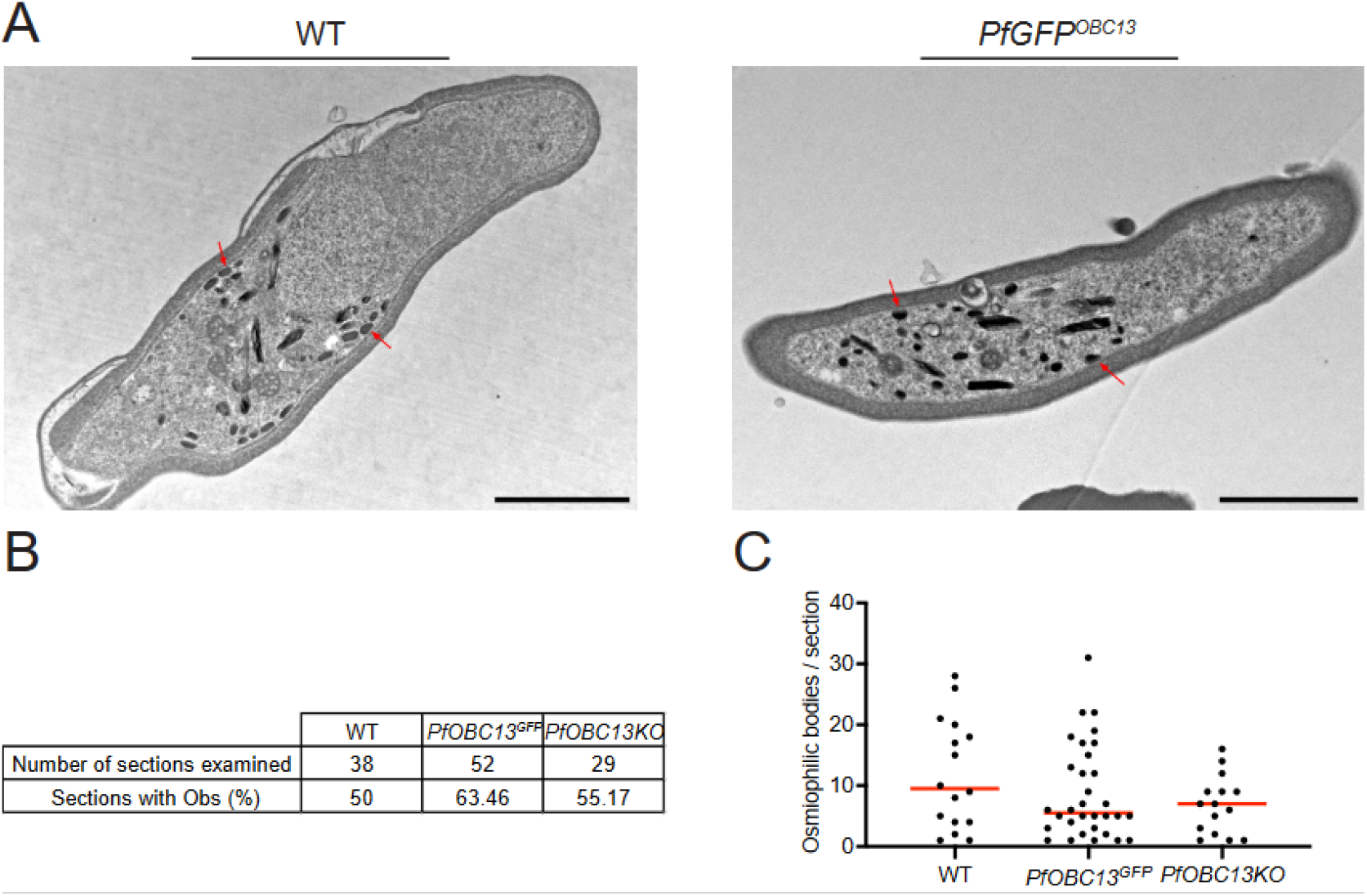
Microscopic analysis of osmiophilic bodies in wild type and transgenic line. The presence and number of osmiophilic bodies was assessed by transmission electron microscopy of wild type (WT), *PfOBC13^GFP^* and *PfOBC13KO* gametocytes. (**A**) Representative images of osmiophilic bodies (red arrows) in the female WT, *PfOBC13^GFP^* and *PfOBC13KO* gametocytes. Scale bar: 2 μm. Prevalence (**B**) and number (**C**, N = 16-33, median shown) of osmiophilic bodies were quantified in sections of WT, *PfOBC13^GFP^* and *PfOB13KO* female gametocytes. Mann-Whitney test was performed to compare samples in **C**. Nonsignificant differences are not indicated.

**Figure S6.**
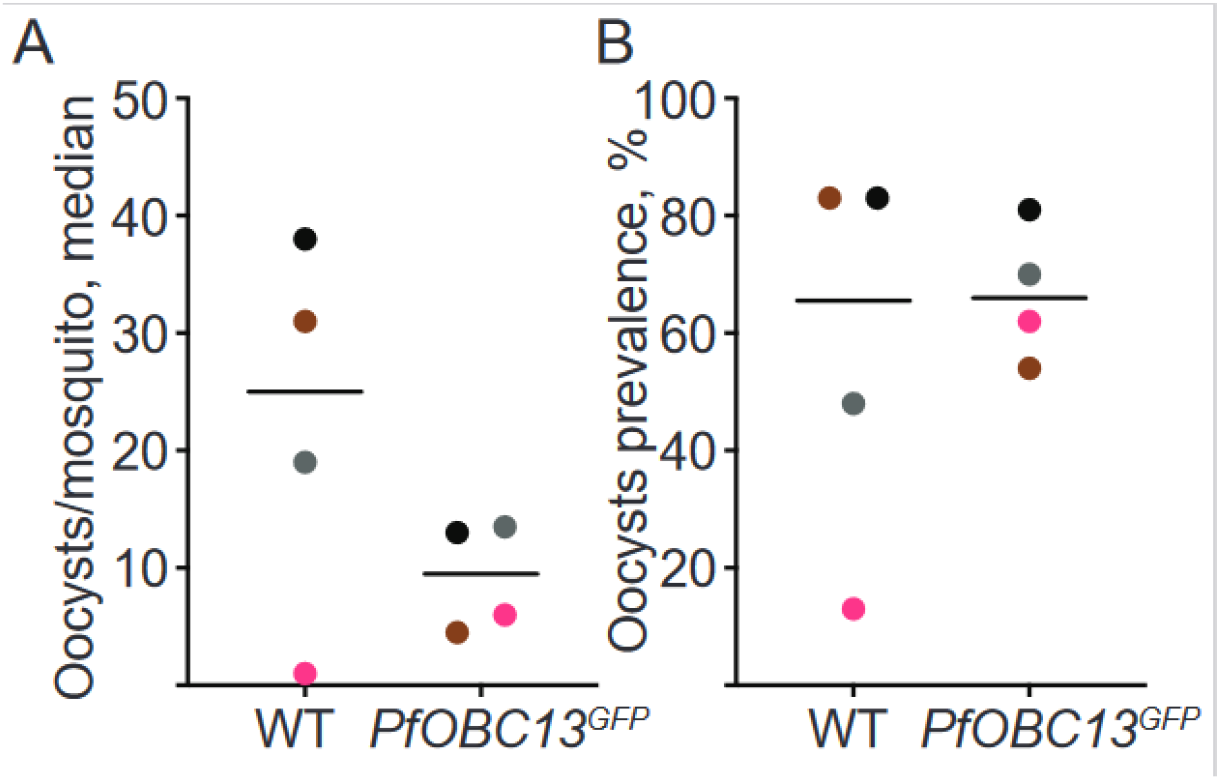
*PfOBC13^GFP^* oocyst development in *A. stephensi* mosquitoes. WT and *PfOBC13^GFP^* oocyst infection intensity (**A,** median of each experiment shown) and prevalence (**B**, %) in *A. stephensi* mosquitoes. Wilcoxon tests were performed to compare samples (N = 4, n = 19-27, median shown). Color code shows paired experiments. Nonsignificant differences are not indicated.

**Figure S7.**
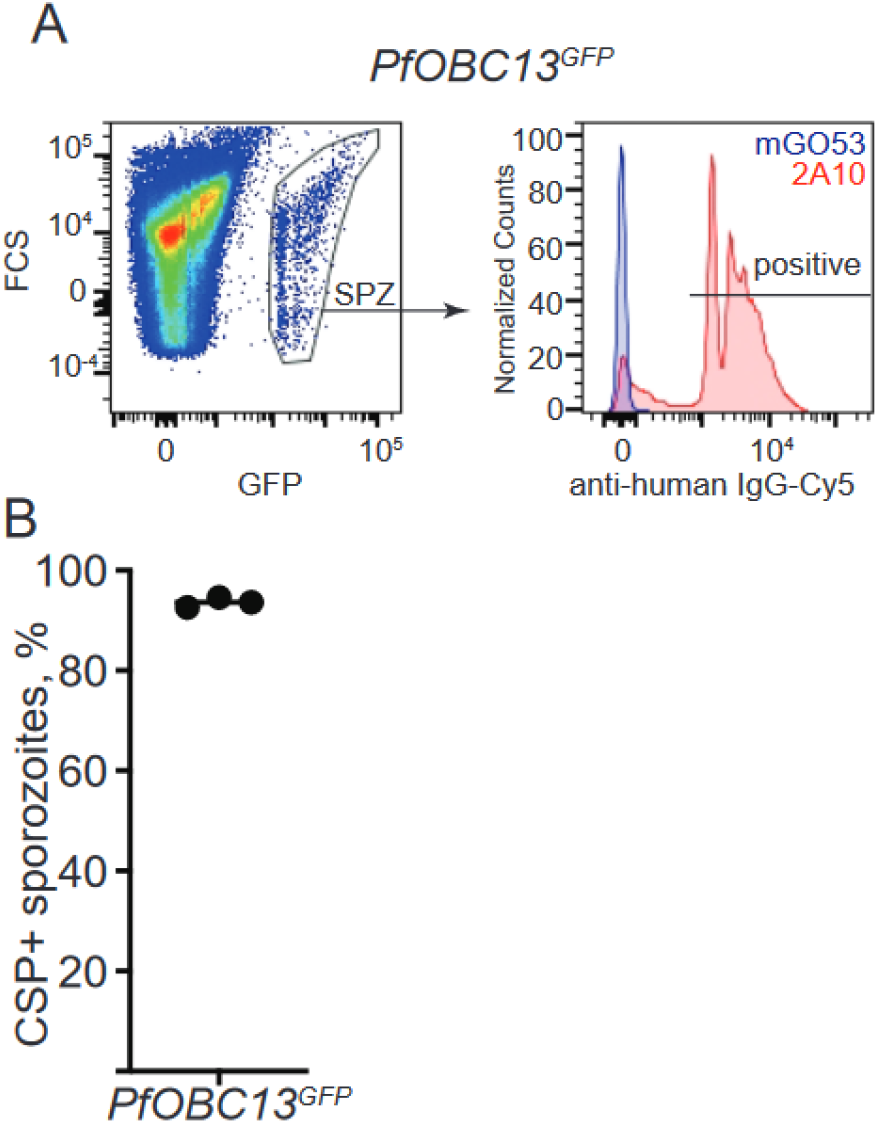
Flow cytometry-based quantification of CSP expression on the surface of *PfOBC13^GFP^*sporozoites. (**A**) Gating strategy for CSP-positive live *PfOBC13^GFP^* sporozoites. The live sporozoite population (SPZ) was gated using the GFP fluorescence signal. The profiles of the anti-CSP mAbs (2A10) and isotype control (mGO53) are shown for comparison. (**B**) Percentage of CSP-positive *PfOBC13^GFP^*sporozoites detected by anti-CSP mAb 2A10 (N = 3, mean shown).

